# Wei2GO: weighted sequence similarity-based protein function prediction

**DOI:** 10.1101/2020.04.24.059501

**Authors:** Maarten J.M.F Reijnders

## Abstract

**Background:** Protein function prediction is an important part of bioinformatics and genomics studies. There are many different predictors available, however most of these are in the form of web-servers instead of open-source locally installable versions. Such local versions are necessary to perform large scale genomics studies due to the presence of limitations imposed by web servers such as queues, prediction speed, and updatability of databases.

**Methods:** This paper describes Wei2GO: a weighted sequence similarity and python-based open-source protein function prediction software. It uses DIAMOND and HMMScan sequence alignment searches against the UniProtKB and Pfam databases respectively, transfers Gene Ontology terms from the reference protein to the query protein, and uses a weighing algorithm to calculate a score for the Gene Ontology annotations.

**Results:** Wei2GO is compared against the Argot2 and Argot2.5 web servers, which use a similar concept, and DeepGOPlus which acts as a reference. Wei2GO shows an increase in performance according to precision and recall curves, Fmax scores, and Smin scores for biological process and molecular function ontologies. Computational time compared to Argot2 and Argot2.5 is decreased from several hours to several minutes.

**Availability:** Wei2GO is written in Python 3, and can be found at https://gitlab.com/mreijnders/Wei2GO

## Introduction

Predicting proteins functions computationally is an integral part of omics research, due to the laborious process of experimentally determining protein functions. This is most apparent in the size difference of Uniprot’s TrEMBL and SwissProt databases, which contain predicted and manually curated proteins respectively [1]. Because of the dependence on predicted protein functions, many software tools have been developed in an attempt to improve the accuracy of electronically annotating these proteins with Gene Ontology (GO) terms [2, 3]. A comprehensive overview of protein function prediction methods is the Critical Assessment of Functional Annotation (CAFA) competition [4]. In its most recent edition, CAFA3, it compared the predictions of 68 participants employing a wide variety of techniques including sequence homology, machine learning, and text mining amongst others. The top-performing predictors in this challenge are considered the state-of-the-art for protein function prediction. With this exposure and proven performance, predictors participating in CAFA have the potential to act as a resource for any user wanting to predict protein functions of their organism of interest. But for CAFA participants to be such a resource, they have two other factors to consider: accessibility and speed. A look at the top performers of CAFA shows that nine out of fourteen teams which placed top ten in one of the GO ontologies, provide their method as a tool accessible through a web server. However, only four out of fourteen teams provide a working locally installable option. Web servers provide an excellent option for the annotation of a limited amount of proteins, but larger amounts are problematic due to computational limitations.

Contrary to web servers, locally installable software provides opportunities to speed up predictions without limitations usually imposed, such as limited protein sequence inputs or queues shared with multiple users. Local versions provide users the means to update the software databases, which is a necessity to provide the most up-to-date and accurate predictions. As a bonus, these versions are often open-source and provide flexibility in the usage of the tool by advanced users, e.g. by utilizing a meta-predictor [5]. As such, there is a need for accurate open-source and locally installable prediction tools.

This paper describes Wei2GO: a weighted sequence similarity and python-based open-source function prediction software. Wei2GO utilizes DIAMOND [6] and HMMER [7] searchers against the UniProtKB [1] and Pfam [8] databases respectively, acquires GO terms through these homology searches, and calculates several weighted scores to accurately associate probabilities to these GO terms. Wei2GO is similar in concept to the web server-based Argot2.5 [9] and its predecessor Argot2 [10], which were top performers in CAFA3 and CAFA2 respectively [4, 11]. Due to this shared niche, this paper provides comparisons between these tools and Wei2GO using precision recall curves, Fmax scores, Smin scores, and a comparison of computational time for their predictions.

## Methods

Wei2GO is a function prediction tool based on sequence similarity with reference proteins. It uses two of the most common sequence alignment algorithms, DIAMOND and HMMER, against the UniProtKB and Pfam database respectively. Each sequence alignment hit is associated with GO terms, taken from the Gene Ontology Annotation (GOA) database [12] for UniProtKB, and pfam2go [13] for Pfam. All GO terms associated with a hit are given a weight relative to their e-value and associated annotation evidence. The weights for each protein-GO term pair are summed across the DIAMOND and HMMER matches. Based on their semantic similarity, the weights of similar GO terms are obtained to get a group score. Finally the GO weight, its information content, and group score are used to calculate the final score associated with the prediction.

### Wei2GO input preparation

Wei2GO requires two input files:

1. Diamond or BLASTP hits against the UniProtKB database in tabular format (-- outfmt 6).
2. HMMScan hits against the Pfam-A database as a table output (--tblout).

These files can either be produced separately, or be incorporated with Wei2GO via its Snakemake pipeline [14].

### Wei2GO algorithm

Each protein matched with a UniProtKB reference via BLAST or DIAMOND is annotated with GO terms associated to these reference proteins in the GOA database. Consecutively, each protein matched with a Pfam domain via HMMScan is annotated with GO terms associated to this domain in the pfam2go database.

For every protein, a GO term *i* is given a weight *W* by taking the sum of the e-value *E* of all UniProtKB and Pfam sequence similarity hits associated with this GO term (1). Terms annotated to a UniprotKB protein with an evidence code other than IEA (Inferred from Electronical Annotation) are given a weight multiplier based on the ratio between the total amount of IEA and non-IEA GO terms present in the GOA database (2).

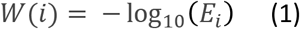

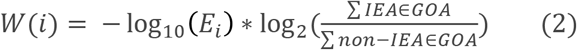

In accordance with the GO Directed Acyclic Graph (DAG) structure, all weights associated to child terms of a GO term are added to its weight.

The total weight associated with each protein-GO term pair is used to calculate the Internal Confidence (InC) score by taking the relative weight of the GO term compared to the weight of the root node (3):

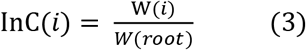

The InC score is used to calculate a Group Score (GS) for each protein-GO term pair, by taking the sum of the InC of all GO terms scoring a semantic similarity of 0.7 or higher with the GO term *j*:

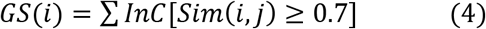

With the semantic similarity (Sim) calculated using Lin’s formula [15] as:

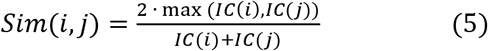

And the Information Content (IC) calculated as a proportion of GO term *i* relative to the size of the entire GOA database:

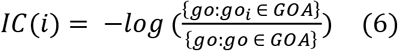

Finally, a total score (TS) for each GO term is calculated as:

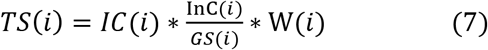

### Test set generation

To assess the performance of Wei2GO it is compared to Argot2.5 and its predecessor Argot2, with the addition of DeepGOPlus [16] which acts as a reference point. All proteins in the GOA database created between 01-01-2018 and 01-06-2020 were extracted. Only GO terms with the evidence code EXP (Inferred from Experiment), IDA (Inferred from Direct Assay), IPI (Inferred from Physical Interaction), IMP (Inferred from Mutant Phenotype), IGI (Inferred from Genetic Interaction), IEP (Inferred from Expression Pattern), TAS (Traceable Author Statement), and IC (Inferred by Curator), were retained. All molecular function terms ‘GO:0005515’ (protein binding) were removed from the annotation set if it was the only molecular function term annotated to the protein, to remove a large amount of bias due to many proteins having this generic term as their only molecular function term. This same approach was used in the CAFA3 test set generation.

The test set was expanded by including all terms according to their GO DAG structure using the ‘is_a’ and ‘part_of’ relationships for parent terms. The final test set consists of 4,193 proteins with 99,453 assigned GO terms of which 1,988 proteins have 61,043 biological process terms, 661 proteins have 4,846 molecular function terms, and 2,466 proteins have 33,564 cellular component terms.

### Test set evaluation

Precision and recall is used for the assessment of Wei2GO. These are calculated as:

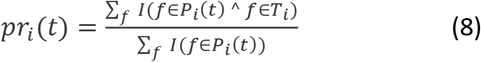

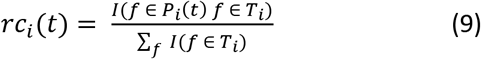

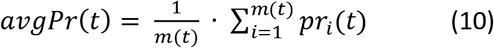

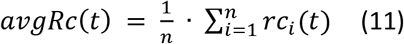

Where for a threshold *t*, the precision *pr* and recall *rc* are calculated per protein *i*. *f* is a GO term, *T* is the set of true GO terms annotated to a protein *i, P* is the set of predicted GO terms annotated to the protein *i*, *m(t)* is the number of proteins with at least one GO term predicted with a score above the threshold, *n* is the total number of proteins, and *I* is an identity function which returns 1 if true and 0 if false.

The optimal harmony between precision and recall is calculated using the maximum F1-score, by calculating the F1-score with 0.01 step-size after transforming the Total Scores to a 0-1 range:

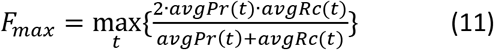

The Smin score is calculated based on the remaining uncertainty *ru* and missing information *mi*, which is the IC of all false negatives and false positives, respectively. A threshold *t* step-size of 0.01 is used after transforming the Total Scores to a 0-1 range.

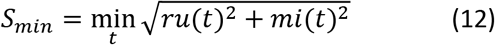

With *ru* as:

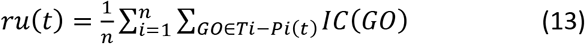

And *mi* as:

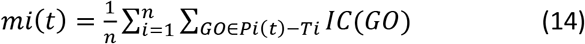

Where *Pi*(t) are all predicted GO terms above a given threshold, and *Ti* are all the GO terms in the true set for a protein.

## Materials

The following datasets and software versions were used:

- UniProt Knowledgebase release 2016-11 for DIAMOND searches
- Pfam-A release 30 for HMMER searches
- DIAMOND version 0.9.11.112
- HMMER version 3.3
- GOA UniProt data release 158 for creating the Wei2GO required data files
- GOA UniProt data release of 13-06-2020 for generating the test set
- go.obo release 02-12-2016
- pfam2go generated based on interpro2go release 31-10-2016
- Argot2 and Argot2.5 were last accessed on 30-10-2020
- DeepGOPlus was downloaded from https://github.com/bio-ontology-research-group/deepgoplus on 30-10-2020.

All datasets used by Wei2GO correspond to the datasets used by Argot2 and Argot2.5.

## Results

Wei2GO is made to facilitate open-source protein function prediction, and the ability to do this independent of web-servers. Two of these web-servers, Argot2.5 and its predecessor Argot2, utilize sequence similarity searches against the UniProtKB and Pfam databases, and apply a weighing algorithm to come to a final prediction score. Wei2GO takes that idea and puts this in an open-source and locally installable concept, which opens up the possibility to update data used by the algorithm independent of the original authors, and the ability to perform protein function prediction on big data.

### Algorithm comparisons between Wei2GO and Argot

Because Wei2GO’s concept is based on that of Argot, it is important to highlight major algorithmic differences between the different software (Table 1). Both Argot’s supplement pfam2go annotations with the GOA GO terms of all proteins belonging to each Pfam entry, which Wei2GO does not. Wei2GO weighs GO annotations differently based on the associated evidence codes in the GOA database, which both Argot’s do not. Argot2.5 is the only software that performs taxonomy-based filtering of GO terms. And finally, Argot2 and Argot2.5 can only handle 5,000 and 10,000 proteins at a time, respectively.

**Table 1:** An overview of the major algorithmic differences between Argot2, Argot2.5, and Wei2GO.

|  | Supplementing<br>pfam2go annotations | Evidence-based<br>weights | Taxonomy-based<br>filtering | Limited number of<br>input sequences |
| --- | --- | --- | --- | --- |
| Argot2 | ✓ | ✗ | ✗ | ✓ |
| Argot2.5 | ✓ | ✗ | ✓ | ✓ |
| Wei2GO | ✗ | ✓ | ✗ | ✗ |

To get an indication of the effect of the algorithmic differences between the software, Table 2 shows an overview of the correctly and incorrectly predicted protein-GO term pairs, irrespective of their prediction scores. The main difference between the software is the amount of false positive (FP) predictions. Argot2, which supplements pfam2go annotations, and to a lesser extend Argot2.5, which supplements pfam2go annotations and applies a taxonomic filter, show a substantially larger amount of FP predictions compared to Wei2GO whilst not showing a large increase in true positive (TP) predictions. Exceptions are Biological Processes (BPO) where Argot2.5 shows both a lower TP and FP count compared to Wei2GO, and Cellular Components (CCO) where Argot2 and Argot2.5 show an increase in TP count. This general trend suggests that supplementing Pfam annotations by adding GOA GO annotations from proteins belonging to the Pfam entry, which is not present in Wei2GO, adds quantity but not quality to the predictions.

**Table 2:** Comparison of Argot2, Argot2.5, and Wei2GO raw prediction counts. All predicted protein-GO term pairs present in the test set are classified as a True Positive (TP) and all predicted protein-GO term pairs not present in the test set are classified as False Positive (FP). Shown are the numbers for Biological Process ontology (BPO), Molecular Function ontology (MFO), and Cellular Component ontology (CCO).

|  | BPO TP | BPO FP | MFO TP | MFO FP | CCO TP | CCO FP |
| --- | --- | --- | --- | --- | --- | --- |
| Argot2 | 40,825 | 2,018,588 | 3,403 | 143,938 | 20,188 | 276,293 |
| Argot2.5 | 14,505 | 109,195 | 3,117 | 174,009 | 14,780 | 53,090 |
| Wei2GO | 38,759 | 124,037 | 3,502 | 37,021 | 11,747 | 35,781 |

### Performance comparisons between Wei2GO and Argot

Several performance metrics are used to compare Wei2GO and Argot: precision-recall curves, Fmax scores, Smin scores, and computational time. DeepGOPlus is added as a part of all these analyses. This deep learning-based and open source protein function prediction software is an excellent example of both predictive performance and usability. Adding this software in the assessments provides perspective as to any performance differences between Wei2GO and Argot. For the purpose of this analysis, all databases used by Wei2GO are the same versions used by Argot2 and Argot2.5.

Precision recall curves were generated for biological process (BPO), molecular function (MFO), and cellular component (CCO) ontologies (Figure 1). This analysis shows that Wei2GO is the best performer in both the BPO and MFO assessment, with most notably BPO having a substantial increase in precision. For CCO, Wei2GO is slightly outperformed by Argot2, while DeepGOPlus performs substantially better than both.

**Figure 1:**
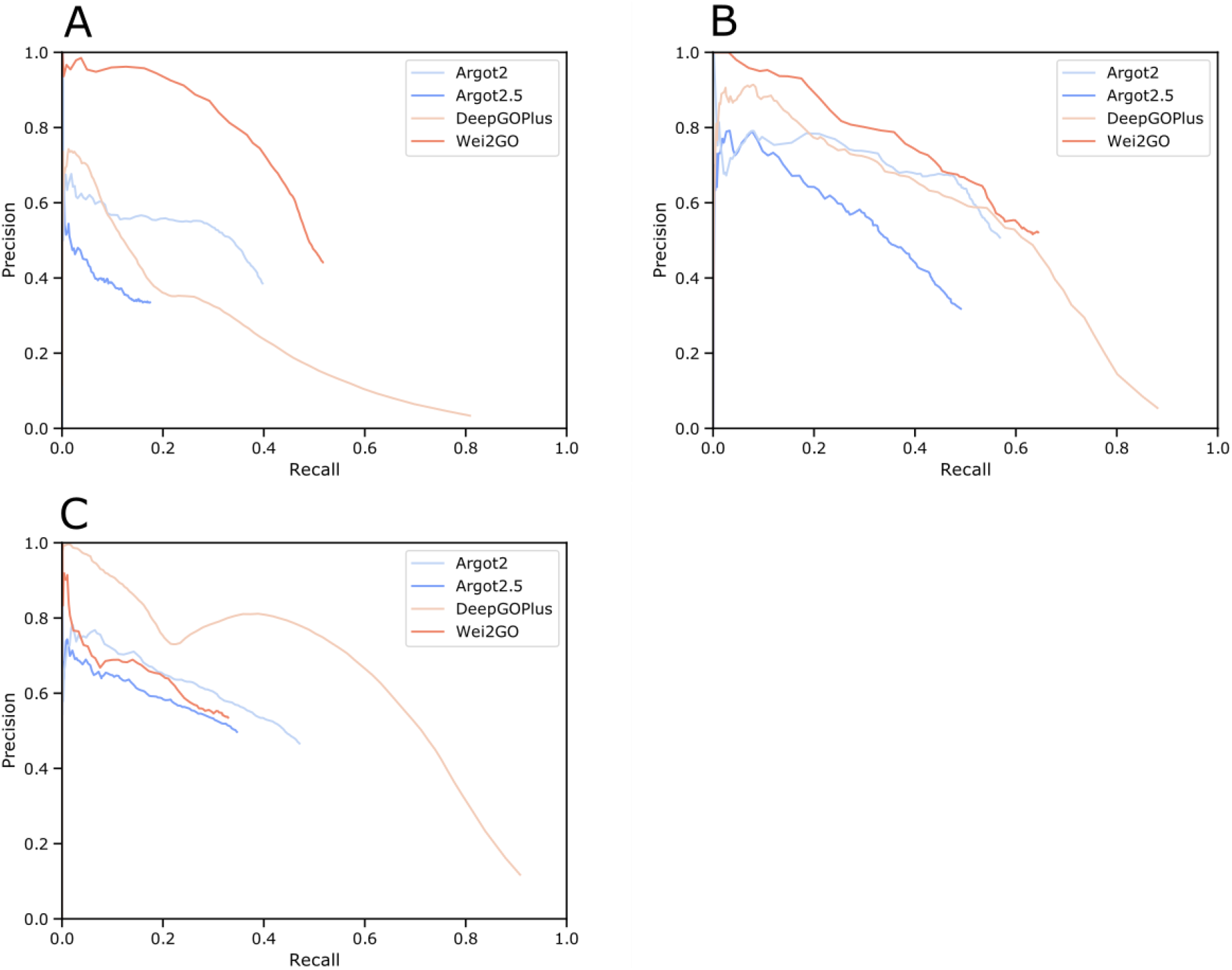
Precision recall curves for (A) biological process, (B) molecular function, and (C) cellular component terms.

Fmax scores were calculated based on the highest F1-score at any given threshold, and Smin scores were calculated based on the lowest S-score at any given threshold (Table 3). The Fmax score is highest in Wei2GO for BPO and MFO, but is outperformed by both Argot2 and DeepGOPlus for CCO. Wei2GO shows the lowest Smin score for both BPO and MFO, and is outperformed by both Argot2 and DeepGOPlus for CCO.

**Table 3:**
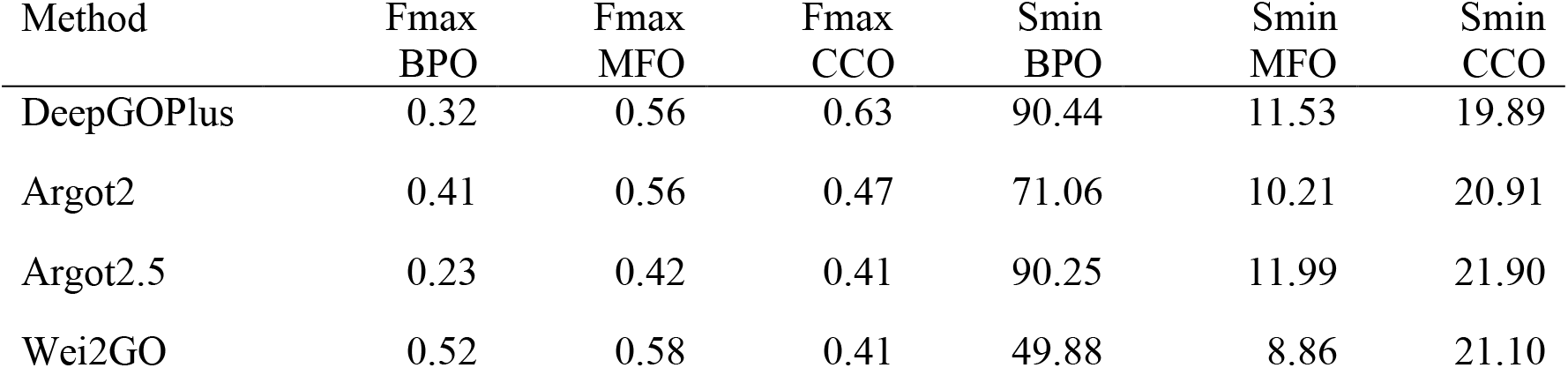
Comparison of the maximum F1-scores (Fmax) and minimum S-scores (Smin) between methods for biological processes (BPO), molecular functions (MFO), and cellular components (CCO).

Computational times were compared in Table 4 based on the 4,193 proteins of the test set. The Argot2 and Argot2.5 web-servers were run without queue times, and all analyses were run on one CPU.

**Table 4:** A comparison of the computational time required for the prediction of the test set used for the analysis in this paper. Based on 4226 proteins. DIAMOND and HMMScan calculations required for the input of Argot2, Argot2.5, and Wei2GO, were excluded from the computational times. No queue times for Argot2 and Argot2.5 were reported.

| Method | Computational<br>time (minutes) |
| --- | --- |
| DeepGOPlus | 2 |
| Argot2 | 192 |
| Argot2.5 | 377 |
| Wei2GO | 4 |

## Discussion

This paper describes Wei2GO: a weighted sequence similarity and python-based open source function prediction software. It utilizes a combination of DIAMOND and HMMER searches, and combines these to extract a probability score associated with GO terms. It is compared against the web-based tools Argot2.5 and Argot2, which share a similar concept, and DeepGOPlus which acts as a reference point for the various metrics used.

Wei2GO shows a large improvement in predicting BPO according to the precision and recall, Fmax, and Smin evaluation metrics. Similarly, MFO shows a less drastic improvement in all comparisons. CCO shows a slight increase in performance over Argot2.5, a slight decrease compared to Argot2, and a substantial decrease in comparison to the reference, DeepGOPlus. All metrics show a reduction in performance for Argot2.5 compared to its predecessor, suggesting taxonomic filtering of GO terms has an adverse effect on prediction performance.

In comparison to the reference, DeepGOPlus, Argot2 similarly shows an improvement in BPO prediction, which implies a sequence similarity-based approach is preferable over a deep learning-based approach. Since Argot2 and Wei2GO share a similar approach and used the same databases for their predictions, performance difference between them implies key algorithmic differences leading to a large improvement in BPO prediction for Wei2GO over Argot2. No definitive conclusion can be made due to the closed-source concept of Argot2, however Table 2 shows Argot2 predicts many more GO terms than Wei2GO whilst not increasing the amount of correct predictions. This is likely due to the former supplementing pfam2go annotations with the GOA annotations of all proteins sharing a Pfam entry.

MFO predictions are similar between Argot2, Wei2GO, and the reference DeepGOPlus. While the same trend is shown in the number of correct and incorrect GO terms predicted by Argot2 and Wei2GO as for BPO (Table 2), the precision and recall curve (Figure 1), Fmax, and Smin scores (Table 3) show Argot2 is better at sorting out the incorrect predictions, leading to a much smaller increase in performance for Wei2GO.

Wei2GO performs worse than Argot2 and DeepGOPlus for CCO predictions. The big difference in precision between DeepGOPlus and the other methods indicates deep learning-based prediction is preferable for this ontology compared to sequence similarity-based prediction. A slight decrease in performance for Wei2GO compared to Argot2 could be due to the relative simple nature of the GO DAG for CCO compared to the other ontologies. Enhancing Pfam domains with the GOA GO terms of its protein members could in such a case improve the predictions.

As observed in the precision recall curves (Figure 1), Wei2GO offers a smoother curve for BPO and MFO predictions compared to Argot2. This is preferable, as it suggests better sorting of high-confidence and low-confidence predictions. While no definitive conclusion can be made due to the closed-source nature of Argot2, this is likely a result of the evidence-based weighing of Wei2GO, as it is more reliable to transfer GO terms through sequence similarity when these come from experimental or curated evidence.

Finally, the computational time for Wei2GO is orders of magnitude lower than that of Argot2 (Table 4), while allowing more proteins to be predicted simultaneously. This opens up the possibility of high-throughput annotating large amounts of omics data with GO terms, as is often needed in the study of comparative genomics and big data studies.

## Conclusion

Wei2GO offers an open-source, locally installable alternative of a proven concept for protein function prediction in the name of Argot2 and Argot2.5. Algorithmic differences in Wei2GO lead to a big performance increase of BPO predictions, while showing similar performance in MFO or CCO predictions. Computational time is decreased from hours down to minutes on a test set of 4,193 proteins, and contrary to Argot, Wei2GO is able to predict an unlimited amount of proteins at a time. In conclusion, Wei2GO not only provides a faster, more flexible, and open-source alternative to Argot2 and Argot2.5, but is able to substantially improve BPO predictions while producing similar performance for MFO and CCO.

### Implementation and availability

Wei2GO requires Python 3.6 or higher, and the NumPy package. Supplementary files used to create the results in this paper, instructions on how to install the software, and instructions on how to run the software can be found on https://gitlab.com/mreijnders/Wei2GO. Additionally, Snakemake scripts are provided to run the full annotation pipeline, and update the databases to their latest versions.

## Acknowledgements

The author would like to thank Dr. Robert M. Waterhouse for offering suggestions and editing parts of the manuscript, and Dr. Bastian V.H. Hornung for proofreading the manuscript.

